# Multi-omics analysis of glucose-mediated signaling by a moonlighting Gβ protein Asc1/RACK1

**DOI:** 10.1101/2021.01.12.426444

**Authors:** Shuang Li, Yuanyuan Li, Blake R. Rushing, Susan L. McRitchie, Janice C. Jones, Susan J. Sumner, Henrik G. Dohlman

**Author notes:** Corresponding authors: Henrik G. Dohlman, Susan J. Sumner.

## Abstract

G proteins were originally discovered through efforts to understand the effects of hormones, such as glucagon and epinephrine, on glucose metabolism. On the other hand, many cellular metabolites, including glucose, serve as ligands for G protein-coupled receptors. Here we investigate the consequences of glucose-mediated receptor signaling, and in particular the role of a Gα subunit Gpa2 and a non-canonical Gβ subunit, known as Asc1 in yeast and RACK1 in animals. Asc1/RACK1 is of particular interest because it has multiple, seemingly unrelated, functions in the cell. The existence of such “moonlighting” operations has complicated the determination of phenotype from genotype. Through a comparative analysis of individual gene deletion mutants, and by integrating transcriptomics and metabolomics measurements, we have determined the relative contributions of the Gα and Gβ protein subunits to glucose-initiated processes in yeast. We determined that Gpa2 is primarily involved in regulating sugar metabolism while Asc1 is primarily involved in amino acid metabolism. Both proteins are involved in regulating purine metabolism. Of the two subunits, Gpa2 regulates a greater number of gene transcripts and was particularly important in determining the amplitude of response to glucose addition. We conclude that the two G protein subunits regulate distinct but complementary processes downstream of the glucose-sensing receptor, as well as processes that lead ultimately to changes in cell growth and metabolism.

**AUTHOR SUMMARY:** Despite the societal importance of glucose fermentation in yeast, the mechanisms by which these cells detect and respond to glucose have remained obscure. Glucose detection requires a cell surface receptor coupled to a G protein that is comprised of two subunits, rather than the more typical heterotrimer: an α subunit Gpa2 and the β subunit Asc1 (or RACK1 in humans). Asc1/RACK1 also serves as a subunit of the ribosome, where it regulates the synthesis of proteins involved in glucose fermentation. This manuscript uses global metabolomics and transcriptomics to demonstrate the distinct roles of each G protein subunit in transmitting the glucose signal. Whereas Gpa2 is primarily involved in the metabolism of sugars, Asc1/RACK1 contributes to production of amino acids necessary for protein synthesis and cell division. These findings reveal the initial steps of glucose signaling and several unique and complementary functions of the G protein subunits. More broadly, the integrated approach used here is likely to guide efforts to determine the topology of complex G protein and metabolic signaling networks in humans.

## INTRODUCTION

All cells respond to changes in extracellular and environmental conditions, many of which are detected by receptors coupled to guanine nucleotide-binding proteins (G proteins). While G protein-coupled receptors (GPCRs) have established roles in detecting odors, light, hormones, and neurotransmitters, more recent investigations have uncovered an important role for GPCRs in responding to nutrients and metabolites such as glucose, amino acids, purine nucleotides, and carboxylic acids including fatty acids (1). GPCR activation leads to the synthesis of chemical second messengers, changes in cell metabolism and transcriptional reprograming. Thus, G proteins act as signal transducers, transmitting a specific extracellular signal to a variety of intracellular second messengers and chemical metabolites. In some cases, the initiating and ensuing signals are one and the same.

The yeast *Saccharomyces cerevisiae* has two G protein signaling systems, one that responds to mating pheromone and another that responds to glucose. These systems do not share components but appear to act in a coordinated fashion; without glucose stimulation, the mating response is delayed until the cells have undergone two complete rounds of cell division (2). Of these systems, the pheromone pathway is the best characterized and is typical of those found in humans. A peptide ligand binds to a cell surface receptor, which then activates a G protein, comprised of an α subunit and a tightly associated βγ subunit dimer. Gα then exchanges GDP for GTP and dissociates from Gβγ. The Gα subunit then activates a phosphatidylinositol 3-kinase while Gβγ initiates a mitogen-activated protein kinase (MAPK) cascade (3, 4). The second GPCR pathway responds to glucose (5). The presumptive glucose receptor (Gpr1) is coupled to a typical Gα (Gpa2) (6–8), but there is no corresponding Gβγ. Rather, Gpa2 appears to assemble with a multifunctional protein called Asc1 (9), known as receptor for activated protein kinase C1 (RACK1) in animals (10). Whereas Gpa2 activates adenylyl cyclase, leading to a transient increase in cellular cAMP (11), Asc1 has the opposite effect on cAMP production (9). It is common in animal cells that Gα and Gβγ bind to and regulate the same effector enzymes, including adenylyl cyclase, in opposition to one another (12).

While Asc1/RACK1 has characteristics of a Gβ subunit, it also has other important functions in the cell. As its name suggests, RACK1 was originally identified as an adaptor for protein kinase C in animals, and was proposed to have a role in kinase-mediated signal transduction (13). RACK1 has also been demonstrated to interact with several GPCRs and G protein βγ subunits (14–17). Most prominently, Asc1/RACK1 is part of the 40S subunit of the ribosome (18–23). In that capacity, Asc1 plays an important role in recruiting quality control systems that diminish frameshifting errors when translation is stalled (24, 25). Moreover, Asc1 regulates a subset of transcripts primarily related to glycolysis, respiration, oxidative stress and fermentation (26). Thus Asc1 is part of two distinct molecular complexes, one involved in glucose sensing and the other in glucose utilization. While the function of Asc1 in the ribosome has been well characterized, its role as an “atypical” G protein is largely unexplored.

Here, we sought to determine the role of Asc1, in comparison with Gpa2, to glucose signaling. To gain a better understanding of their relative contributions to cell physiology, we undertook an integrated metabolomics and transcriptomics analysis, comparing mutants that lack the Gα or Gβ subunit. By this approach, one that is largely unprecedented in the GPCR field, we have identified the earliest events leading to glucose fermentation. Our analysis is likely to guide similar efforts to determine the topology of complex G protein and metabolic signaling networks in humans

## RESULTS

We determined previously that Asc1 binds to Gpa2, that these proteins have opposing effects on adenylyl cyclase activity, and that they act in response to the glucose receptor Gpr1 (9). Here we sought a comprehensive understanding of the molecular and cellular consequences of G protein activation. To that end, we undertook a multi-platform investigation, performing untargeted metabolomics and transcriptomics analysis, in cells lacking each of these proteins, in response to glucose. By measuring changes in gene expression and perturbations in host metabolism we sought to gain an understanding of Asc1, apart from its alternative role in translation, and how it complements the functions of Gpa2.

Wildtype, *gpr1, gpa2* and *asc1* cells (all without nutritional auxotrophic markers) were grown for 1 h in low (L, 0.05%) glucose, then divided and either left untreated or treated with high (H, 2%) glucose for 2 min (metabolomics) or 10 min (transcriptomics). These time points were selected based on prior data, showing an early and transient spike of cAMP and a subsequent induction of genes within 10 minutes of glucose treatment (see Methods) (27). We then analyzed our data using Principal Component Analysis (PCA). This unsupervised multivariate analysis method is particularly useful for the visualization of the relationship between observations and variables. When applied to our transcriptomics data, PCA indicated good differentiation of groups based on the proximity of data points for a given treatment and genotype (Figure 1A). This analysis revealed that PC1, which aligns primarily with treatment, accounts for 90% of variance while PC2, which aligns primarily with genotype, represents 5% of variance. Thus the first 2 components explained 95% of the variance. For metabolomics, the first 2 components explained 63% of the variance (Figure 1B). For both measurements, and as expected, *gpr1* aligned closely with *gpa2* (6–9, 11). Both measures are consistent with previously established opposing effects of Gα and Gβ on processes downstream of the G protein.

**Figure 1.**
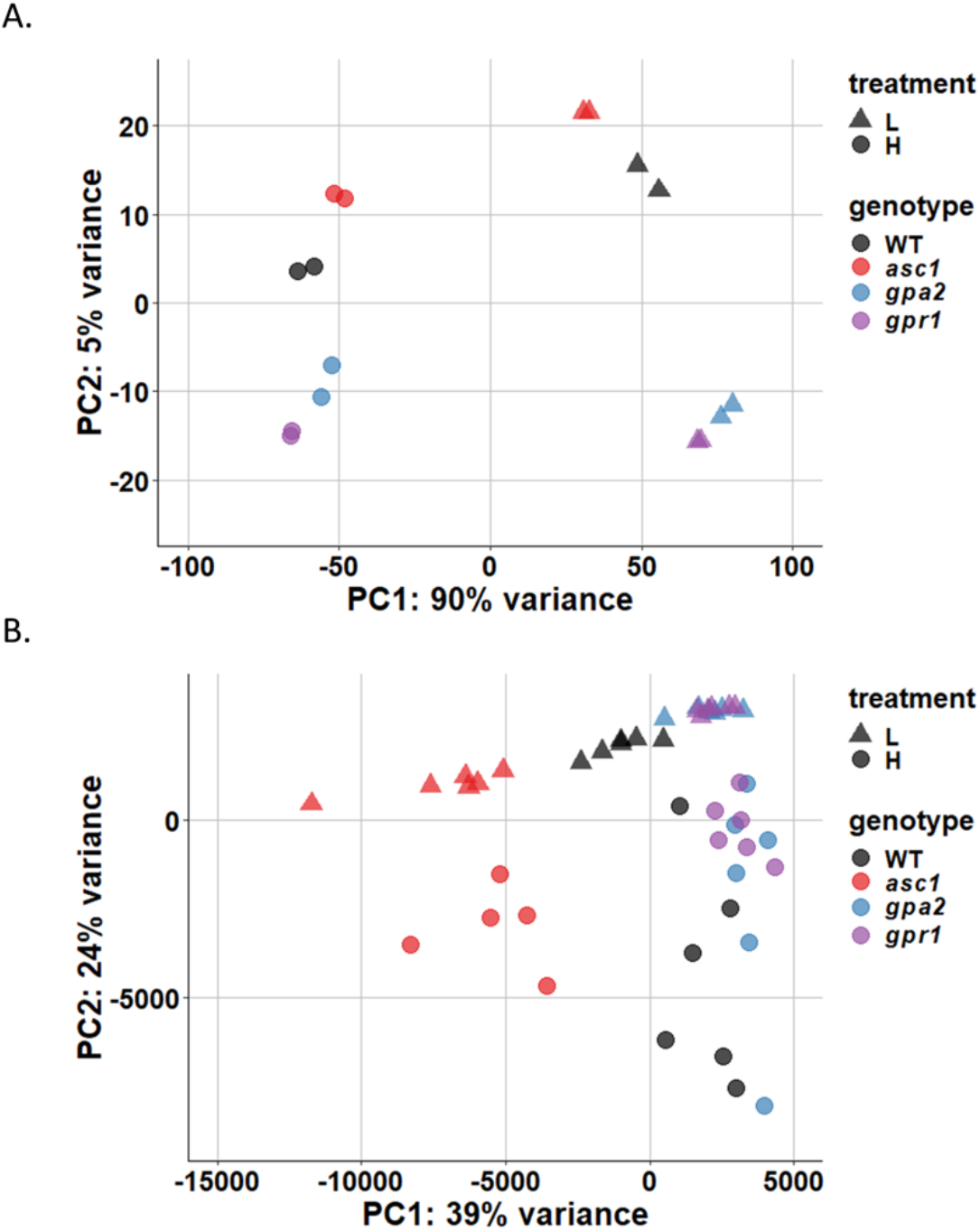
PCA plots. For A) transcriptomics and B) metabolomics, X-axis shows PC1 with the percentage of explained variance and Y-axis shows PC2 with the percentage of explained variance. Data are scaled as detailed in Methods. Wildtype (black), *asc1* (red), *gpa2* (blue), *gpr1* (purple). Low glucose (L, 0.05% glucose)-triangles, high glucose (H, 2% glucose)-circles.

Our next objective was to identify the specific pathways and processes regulated by each G protein subunit. To that end, we analyzed the transcriptomics and metabolomics data, with and without glucose addition, for wildtype, *gpr1, gpa2* and *asc1* cells. Herein we use the term “concentration analysis” when comparing the data for mutant and wildtype cells after the addition of high glucose (Figures 2A and 2C), and we use the term “sensitivity analysis” when comparing the difference in response at high and low glucose for the mutants (mutantH-mutantL) and the wildtype (wtH-wtL) cells (Figures 2B and 2C). Concentration analysis reveals cell status after glucose addition, while sensitivity analysis reveals response amplitude (H-L) differences for the different strains. Such amplitude differences are important when a negative regulator restores the system to baseline in the face of sustained (or step-wise) activation by a positive regulator.

**Figure 2.**
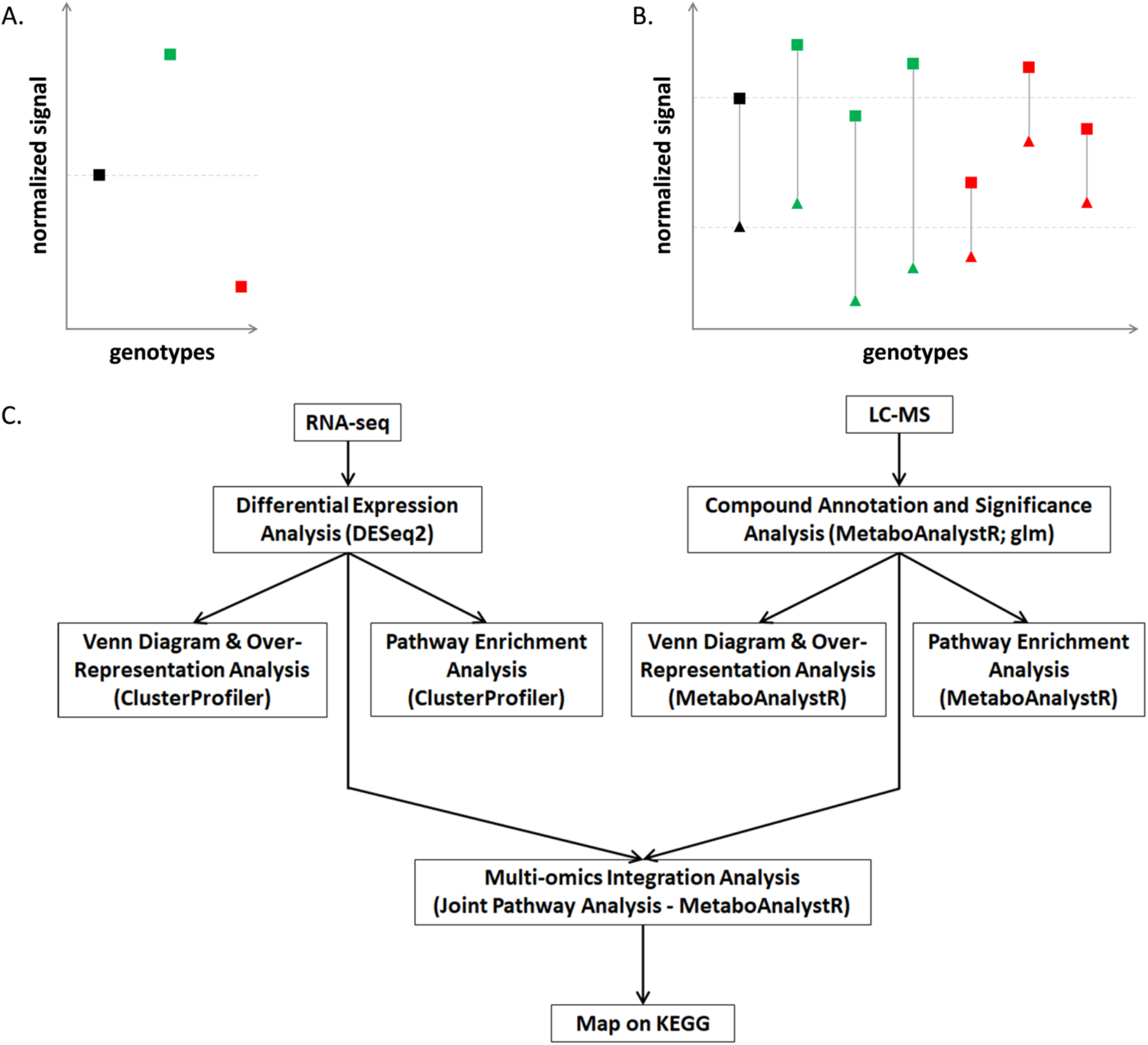
Concentration and sensitivity analysis. Illustration of different modes of changes as captured by concentration analysis (A) and sensitivity analysis (B). The Y-axis represents either normalized gene counts or normalized peak area for metabolites. The X-axis represents different genotypes. A hypothetical wildtype is shown in black. The triangle represents measurement at low glucose and the square represents measurement at high glucose. The connecting grey line represents the response amplitude, detected by sensitivity analysis (wtH-wtL). Hypothetical mutants with increased response amplitude are colored green, while mutants with decreased response amplitude are colored red. C) analysis pipeline of concentration analysis and sensitivity analysis for transcriptomics and metabolomics data.

In summary, we present integrated transcription data and untargeted metabolomics data obtained for the cell extracts, as follows:

Concentration Analysis

A. *gpa2* mutant vs wildtype at high (2%) glucose
B. *asc1* mutant vs wildtype at high (2%) glucose

We then compare the results of A and B, calculated as detailed in Methods.

Sensitivity Analysis

A. difference between *gpa2* mutant at high and low glucose (gpa2H-gpa2L) vs. difference between wildtype at high and low glucose (wtH-wtL)
B. difference between *asc1* mutant at high and low glucose (asc1H-asc1L) vs. difference between wildtype at high and low glucose (wtH-wtL)

We then compare the results of A and B, calculated as detailed in Methods.

### Concentration analysis

We began by establishing the transcriptional and metabolic profile of wildtype cells. We defined the differentially-expressed genes (DEGs) as having an adjusted p-value <0.05 and absolute fold-change value >1. Then, using the ClusterProfiler package in R (28) we performed gene set enrichment analysis (GSEA) in the Kyoto Encyclopedia of Genes and Genomes (KEGG) (29–31). This database provides an overview of biological pathways in the cell, as determined by genome sequencing and other high-throughput methods. GSEA determines whether a defined set of genes shows statistically significant and concordant differences between two phenotypes. When comparing wildtype before and after glucose addition (0.05% vs. 2%), we observed transcriptomic changes for 2500 DEGs in 32 pathways (adjusted p-value <0.05), including ribosome, DNA replication, transcription, cell cycle as well as carbon, amino acids, lipids and nucleotide metabolism (Table 1). Note that the same p-value threshold was used throughout. These differences were expected, and reflect processes needed to transition from a low glucose phase where metabolism supports cellular homeostasis (e.g. autophagy, addressing reactive oxygen species, maintaining osmotic balance) to a high glucose phase where metabolism supports cell growth and division (structural rearrangements as well as anabolic processes to make building blocks that support cell proliferation) (32). We next conducted untargeted metabolomics by mass spectrometry, as detailed in Methods. Pathway enrichment analysis was performed in MetaboAnalystR (31,32), using the Fisher’s method to integrate Mummichog (33) and GSEA results to produce the combined p-values reported here (Table 1). The pathway enrichment analysis module Mummichog is optimized for detecting prominent changes while GSEA excels at detecting concordant small changes in peak intensity. Because of the uncertainty associated with peak annotation for LC-MS data, the reliability of pathway enrichment is improved when combining the results from two different statistical methods (31,32). When comparing 0.05% to 2% glucose in wildtype, 11 pathways were perturbed with a combined p-value <0.05 (same threshold throughout) (Table 1). These include perturbations in the metabolism of sugars, amino acids, nucleotides and lipids, and are concordant with changes in gene transcription. Again, these differences reflect processes needed to prepare the cell for growth and division. In addition, they are likely related to the role of Asc1 and Gpa2 in haploid invasive growth (9, 33, 34), a process where cells form long branching filaments and exhibit increased adherence and invasion of the substratum during periods of glucose limitation (35).

**Table 1.** Single-omics analysis results for wildtype between 2% (H) and 0.05% (L) glucose. First block shows GSEA for transcriptomics with adjusted p-value <0.05, arranged in ascending order; second block shows MetaboAnalystR pathway enrichment analysis for metabolomics with combined p-value <0.05 arranged in ascending order, as detailed in Methods.

| Transcriptomics |  |  |  | Metabolomics |  |
| --- | --- | --- | --- | --- | --- |
| enriched pathways | adjusted p-value | enriched pathways | adjusted p-value | enriched pathways | combined p-value |
| Ribosome | 0.0023 | Lysine biosynthesis | 0.0051 | Starch and sucrose metabolism | 0.0010 |
| Cell cycle - yeast | 0.0023 | Fructose and mannose metabolism | 0.0075 | Tyrosine metabolism | 0.0065 |
| Biosynthesis of amino acids | 0.0023 | Aminoacyl-tRNA biosynthesis | 0.0080 | Galactose metabolism | 0.0073 |
| RNA transport | 0.0023 | Sulfur metabolism | 0.0094 | Cysteine and methionine metabolism | 0.0184 |
| Ribosome biogenesis in eukaryotes | 0.0023 | One carbon pool by folate | 0.0098 | Glycolysis / Gluconeogenesis | 0.0198 |
| Cysteine and methionine metabolism | 0.0023 | Valine, leucine and isoleucine biosynthesis | 0.0101 | Inositol phosphate metabolism | 0.0207 |
| RNA polymerase | 0.0023 | Proteasome | 0.0101 | Amino sugar and nucleotide sugar metabolism | 0.0233 |
| Purine metabolism | 0.0028 | Autophagy - yeast | 0.0184 | Purine metabolism | 0.0343 |
| DNA replication | 0.0028 | Pyrimidine metabolism | 0.0184 | N-Glycan biosynthesis | 0.0361 |
| Galactose metabolism | 0.0028 | Amino sugar and nucleotide sugar metabolism | 0.0188 | Pentose phosphate pathway | 0.0364 |
| Glyoxylate and dicarboxylate metabolism | 0.0028 | Selenocompound metabolism | 0.0226 | Butanoate metabolism | 0.0443 |
| Citrate cycle (TCA cycle) | 0.0028 | Mismatch repair | 0.0231 |  |  |
| Peroxisome | 0.0028 | Glycine, serine and threonine metabolism | 0.0322 |  |  |
| Starch and sucrose metabolism | 0.0028 | Glycolysis / Gluconeogenesis | 0.0387 |  |  |
| Oxidative phosphorylation | 0.0033 | Autophagy - other | 0.0416 |  |  |
| Carbon metabolism | 0.0041 | Meiosis - yeast | 0.0474 |  |  |

We next considered the effects of glucose in each of the mutant strains and how each mutant compares with wildtype. When measuring transcript abundance after the addition of 2% glucose, we found 277 and 170 DEGs for *gpa2* vs. wildtype and for *asc1* vs. wildtype, respectively. The *gpa2* mutant exhibited changes in transcripts linked to oxidative phosphorylation and ribosome biogenesis (Table 2 and Table S1, Supporting Information). The *asc1* mutant was enriched for twelve pathways including ribosome biogenesis and the biosynthesis of amino acids (Table 3 and Table S2, Supporting Information). Figure 3A depicts a Venn diagram, comparing the DEGs for *gpa1* vs. wildtype and for *asc1* vs. wildtype (Table S3, Supporting Information). These comparisons show that *gpa2* and *asc1* had similar effects on 28 DEGs, of which 21 were up-regulated and seven were down-regulated. The *asc1* mutant uniquely affected 117 DEGs (90 up-regulated, 27 down-regulated) and the *gpa2* mutant uniquely affected 224 DEGs (94 up-regulated, 130 down-regulated). Gpa2 and Asc1 both targeted an additional 25 DEGs with opposing effects on their expression, as compared with wildtype. Most of the opposing effects were related to retrotransposon elements and are unlikely to be related to glucose metabolism or signaling. We also conducted over-representation analysis (ORA) for the unique as well as shared intersects of the diagram. The shared DEGs were over-represented for arginine biosynthesis (Figure 3B). Arginine contributes to nitrogen balance, through urea production, and also regulates protein translation (36). In comparison the unique DEGs were over-represented for eight pathways related mainly to amino acid metabolism (*asc1*, Figure 3C) and six pathways linked to glucose and energy metabolism (*gpa2*, Figure 3D). Thus, pathway analysis revealed that Asc1 has effects that are broader than those of Gpa2.

**Table 2.** Single- and Multi-omics integration results for gpa2 by concentration analysis. First block shows GSEA for transcriptomics with adjusted p-value <0.05, arranged in ascending order; second block shows MetaboAnalystR pathway enrichment analysis for metabolomics with combined p-value <0.05, arranged in ascending order; third block shows MetaboAnalystR joint pathway analysis with adjusted p-value <0.05, arranged in ascending order, as detailed in Methods.

| Transcriptomics |  | Metabolomics |  | Integration |  |
| --- | --- | --- | --- | --- | --- |
| enriched pathways | adjusted p-value | enriched pathways | combined p-value | enriched pathways | adjusted p-value |
| Ribosome biogenesis in eukaryotes | 0.0084 | Purine metabolism | 0.0021 | Oxidative phosphorylation | 3.13E-14 |
| Oxidative phosphorylation | 0.0084 | Fructose and mannose metabolism | 0.0047 | Galactose metabolism | 1.60E-11 |
|  |  | Amino sugar and nucleotide sugar metabolism | 0.0079 | ABC transporters | 1.31E-08 |
|  |  | Galactose metabolism | 0.0079 | Glycolysis or Gluconeogenesis | 8.02E-05 |
|  |  | Glutathione metabolism | 0.0155 | Fructose and mannose metabolism | 8.18E-05 |
|  |  | Tyrosine metabolism | 0.0224 | Starch and sucrose metabolism | 8.18E-05 |
|  |  | Arginine biosynthesis | 0.0234 | Arginine biosynthesis | 0.0012 |
|  |  | Biotin metabolism | 0.0252 | Pentose phosphate pathway | 0.0012 |
|  |  | Aminoacyl-tRNA biosynthesis | 0.0360 | Purine metabolism | 0.0017 |
|  |  | Starch and sucrose metabolism | 0.0396 | Amino sugar and nucleotide sugar metabolism | 0.0073 |
|  |  | Phosphatidylinositol signaling system | 0.0424 | beta-Alanine metabolism | 0.0119 |
|  |  |  |  | Alanine, aspartate and glutamate metabolism | 0.0139 |
|  |  |  |  | Cysteine and methionine metabolism | 0.0149 |
|  |  |  |  | Citrate cycle (TCA cycle) | 0.0221 |

**Table 3.** Single- and Multi-omics integration results for *asc1* by concentration analysis. First block shows GSEA for transcriptomics with adjusted p-value <0.05, arranged in ascending order; second block shows MetaboAnalystR pathway enrichment analysis for metabolomics with combined p-value <0.05, arranged in ascending order; third block shows MetaboAnalystR joint pathway analysis with adjusted p-value <0.05, arranged in ascending order, as detailed in Methods.

| Transcriptomics |  | Metabolomics |  | Integration |  |
| --- | --- | --- | --- | --- | --- |
| enriched pathways | adjusted p-value | enriched pathways | combined p-value | enriched pathways | adjusted p-value |
| Starch and sucrose metabolism | 0.0155 | Arginine biosynthesis | 0.0029 | Arginine biosynthesis | 7.47E-10 |
| Biosynthesis of amino acids | 0.0156 | Glutathione metabolism | 0.0054 | ABC transporters | 7.47E-10 |
| Sulfur metabolism | 0.0156 | Aminoacyl-tRNA biosynthesis | 0.0214 | Purine metabolism | 1.41E-09 |
| Ribosome | 0.0156 | Purine metabolism | 0.0481 | Tryptophan metabolism | 0.0074 |
| Galactose metabolism | 0.0156 |  |  | Galactose metabolism | 0.0074 |
| Meiosis - yeast | 0.0156 |  |  | Cysteine and methionine metabolism | 0.0087 |
| Glycine, serine and threonine metabolism | 0.0243 |  |  | Alanine, aspartate and glutamate metabolism | 0.0087 |
| One carbon pool by folate | 0.0243 |  |  | Histidine metabolism | 0.0087 |
| Aminoacyl-tRNA biosynthesis | 0.0243 |  |  | Arginine and proline metabolism | 0.0136 |
| 2-Oxocarboxylic acid metabolism | 0.0387 |  |  | Phenylalanine metabolism | 0.0191 |
| Lysine biosynthesis | 0.0477 |  |  | Lysine biosynthesis | 0.0348 |
| Ribosome biogenesis in eukaryotes | 0.0477 |  |  | Amino sugar and nucleotide sugar metabolism | 0.0465 |

**Figure 3.**
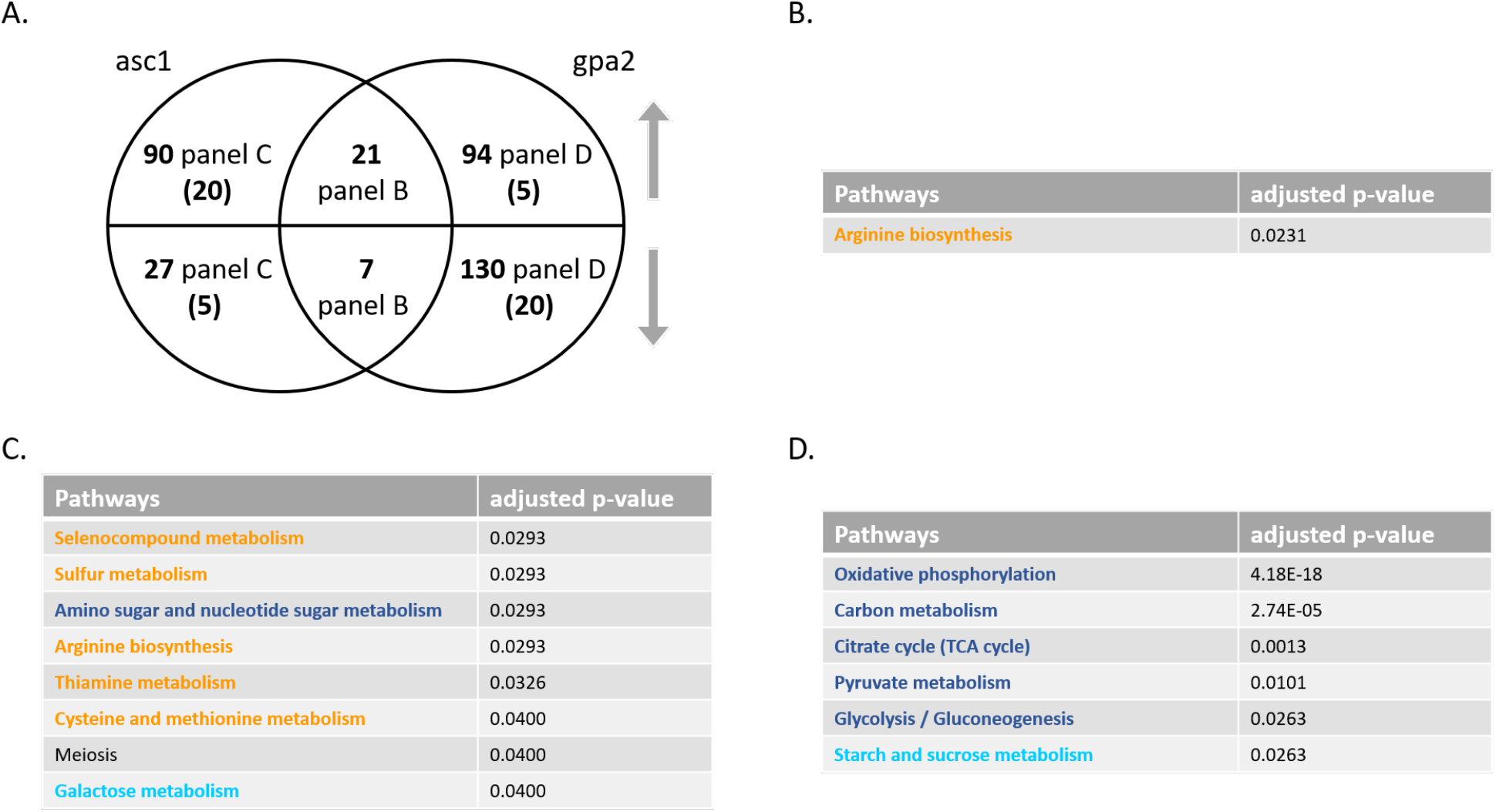
Concentration analysis of differentially expressed genes (DEGs) after glucose addition. A) Venn diagram of subsets of DEGs, for *asc1* and *gpa2* vs. wildtype, after glucose addition to 2%. Upper semicircle shows up-regulated DEGs and lower semicircle shows down-regulated DEGs. Numbers in parenthesis are shared DEGs regulated in the opposite direction, placed in the area corresponding to the direction of regulation. DEGs used for ORA analysis that are B) shared and change in the same direction; C) unique to *asc1*; D) unique to *gpa2*. Pathway names are color coded according to the KEGG metabolic pathway map: glucose and ATP (blue), starches and non-glucose sugars (cyan), amino acids (yellow), nucleotides (red), lipids and glycerophospholipids (teal), cofactors and vitamins (pink). All-gene pathways not included in the KEGG metabolic pathway map are in black. All pathways with adjusted p-value <0.05 are listed, in ascending order.

We next conducted untargeted metabolomics, followed by metabolite identification/annotations and pathway enrichment analysis, for *gpa2* and *asc1* (Tables S1 and S2 respectively, Supporting Information). When compared to wildtype, *gpa2* cells were enriched in ten pathways including fructose and mannose as well as purine metabolism (Table 2), while *asc1* cells were enriched in four pathways including arginine biosynthesis and purine metabolism (Table 3). Peak annotations derived from MetaboAnalystR are hereafter referred to as “metabolites.”. The Venn diagram shows shared and unique significantly perturbed metabolites (SPMs, defined as those with p<0.05) for each mutant vs wildtype comparison (Figure 4A and Table S4, Supporting Information). In this analysis, *gpa2* and *asc1* had similar perturbations for 28 SPMs, 23 of which were increased and five of which were decreased in both mutants. The *asc1* mutation uniquely perturbed an additional 47 SPMs (44 increased, 4 decreased) compared to wildtype, while *gpa2* uniquely affected an additional 33 SPMs (24 increased, 9 decreased). When comparing the two mutants, ORA revealed that those SPMs that changed in the same direction were enriched in pathways related to sugar metabolism: amino sugar and nucleotide sugar metabolism, as well as fructose and mannose metabolism (Figure 4B). Because nucleotide sugars (e.g. UDP-glucose) are substrates for protein and lipid glycosylation, they may reflect preparation for new cell wall synthesis (37). Glycosylation of the mucin Msb2 is needed to ensure signal fidelity downstream of the filamentous/invasive growth pathway (38). SPMs perturbed in the opposite direction were enriched in purine metabolism (Figure 4C). The perturbations unique to individual mutants were over-represented for two pathways related to amino acid metabolism (*asc1*, Figure 4D) and two pathways linked to non-glucose sugar metabolism (*gpa2*, Figure 4E). Contrary to what we observed for transcriptomics, pathway enrichment analysis for metabolomics revealed that the *gpa2* mutant had an impact that was broader than that of the *asc1* mutant (Tables 2 and 3). ORA analysis with subsets of SPMs corroborated the trends observed in transcriptomics. Based on these data we conclude that Asc1 mainly affects metabolites related to amino acids, Gpa2 mainly affects sugar metabolism, and the two proteins have opposing effects on purine metabolism.

To gain a deeper understanding of the functional relationship between changes in gene transcription and changes in the levels of metabolites, we employed the joint pathway analysis module in MetaboAnalystR. In this application, we input all significantly perturbed genes (DEGs) and significantly perturbed metabolites (SPMs), and queried for those over-represented in KEGG. By integrating the data in this manner, we sought to obtain more information than could be gleaned from transcriptomics and metabolomics separately. We found that, when compared to wildtype cells, *gpa2* mutants were enriched in fourteen pathways while *asc1* mutants were enriched in twelve pathways (Tables 2 and 3, adjusted p-value <0.05 throughout). Both mutants affected genes or metabolites involved in the synthesis of amino acids (cysteine, methionine, arginine, alanine, aspartic acid and glutamic acid), the metabolism of purines, galactose, amino sugars and nucleotide sugars, as well as genes/metabolites involved in ABC transporters. These results reveal a shared role of Asc1 and Gpa2 in regulating the metabolism of sugars, as expected for any component of the glucose-sensing pathway. In addition, both subunits affected the metabolism of amino acids, particularly branches of that pathway most closely linked to the TCA cycle. Amino acids can be used for energy production in the TCA cycle, which is common when glucose is being syphoned off for anabolic processes such as making nucleotides via the pentose phosphate pathway (Figures S1-6, Supporting Information). The *gpa2* strain was unique in regulating gluconeogenesis, oxidative phosphorylation, TCA cycle, and the pentose phosphate pathway as well as the metabolism of β-alanine, starch, sucrose, fructose and mannose (Table 2). The pentose phosphate pathway is an offshoot of glycolysis that supplies de novo nucleotide biosynthesis. Thus, it appears that the Gα subunit Gpa2 regulates the conversion of glucose to ATP and molecular building blocks such as lipids and amino acids. In contrast, the Gβ subunit Asc1 is unique in regulating a variety of amino acids (Table 3). Notably, most of the changes observed for *gpa2* and *asc1* were also observed in cells lacking their shared activator, the GPCR Gpr1 (Table S5, Supporting Information). Thus, it appears that the Gβ subunit specifically regulates the utilization of nitrogen in part through the metabolism of arginine and other amino acids. More broadly, the two G protein subunits regulate distinct but complementary processes downstream of the glucose sensing receptor.

**Figure 4.**
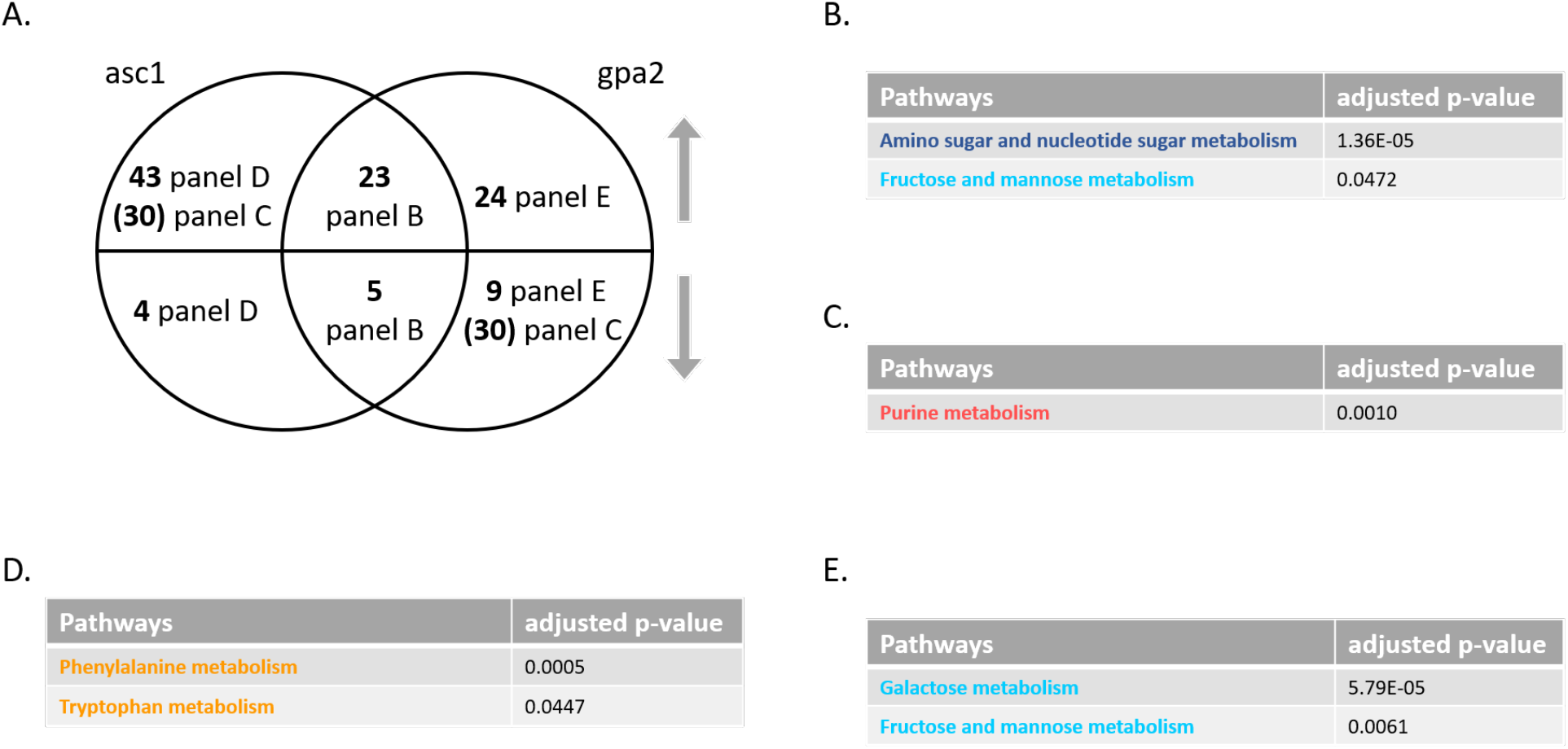
Concentration analysis of significantly perturbed metabolites (SPMs) after glucose addition. A) Venn diagram of subsets of SPMs, for *asc1* and *gpa2* vs. wildtype, after glucose addition. Upper semicircle shows up-regulated SPMs and lower semicircle shows down-regulated SPMs. Numbers in parenthesis are shared SPMs regulated in the opposite direction, placed in the area corresponding to the direction of regulation. SPMs used for ORA analysis that are B) shared and change in the same direction; C) shared and change in the opposite direction; D) unique to *asc1*; E) unique to *gpa2*. Pathway names are color coded according to the KEGG metabolic pathway map as detailed in Figure 3. All pathways with adjusted p-value <0.05 are listed, in ascending order.

To visualize the functional relationship of Gpa2 and Asc1, we projected the inputs of our integration analysis onto the comprehensive yeast Metabolic Pathways diagram provided in KEGG. The map is color coded to delineate glucose and ATP (blue), starches and non-glucose sugars (cyan), amino acids (yellow), nucleotides (red), lipids and glycerophospholipids (teal), cofactors and vitamins (pink) (Figure S7, Supporting Information). More specifically, we highlighted the products of genes (black lines) and metabolites (black dots) that were perturbed and deemed to be statistically significant (DEGs and SPMs), and used gray boxes to delineate clusters associated with an enriched pathway. When comparing clusters it was evident that changes in the *gpa2* strain compared to wild type were concentrated in regions related to glucose and ATP as well as starch and non-glucose sugars (Figure 5), while the changes in *asc1* were concentrated in various types of amino acid metabolism (Figure 6). Both mutants also impacted purine metabolism, but they did so in opposition to one another as detailed above (Figure 4C). These changes mirror the opposing effects of Gpa2 and Asc1 on cAMP. More broadly, these findings highlight the differences between the two G protein subunits and their effects downstream of the glucose-sensing receptor. Whereas Gα regulates sugar utilization, Gβ regulates amino acids.

MetaboAnalystR is well suited for annotating a large number of signals. A complementary approach is to use our in-house library annotation, which includes retention time (RT), exact mass, and MS/MS library (OL) developed with data acquired for standards run under the same conditions as the study samples, as well as matching to public databases (PD), as described in Supporting Information. Metabolites are reported (Table S6, Supporting Information) based on the confidence in the assignment (e.g. OL1 is a match to the in-house library by retention time, exact mass, and MS/MS fragmentation; PDa is matched to a public database by mass and experimental MS/MS). The signals identified and annotated by this method yielded pathways that mirrored those obtained using MetaboAnalystR.

**Figure 5.**
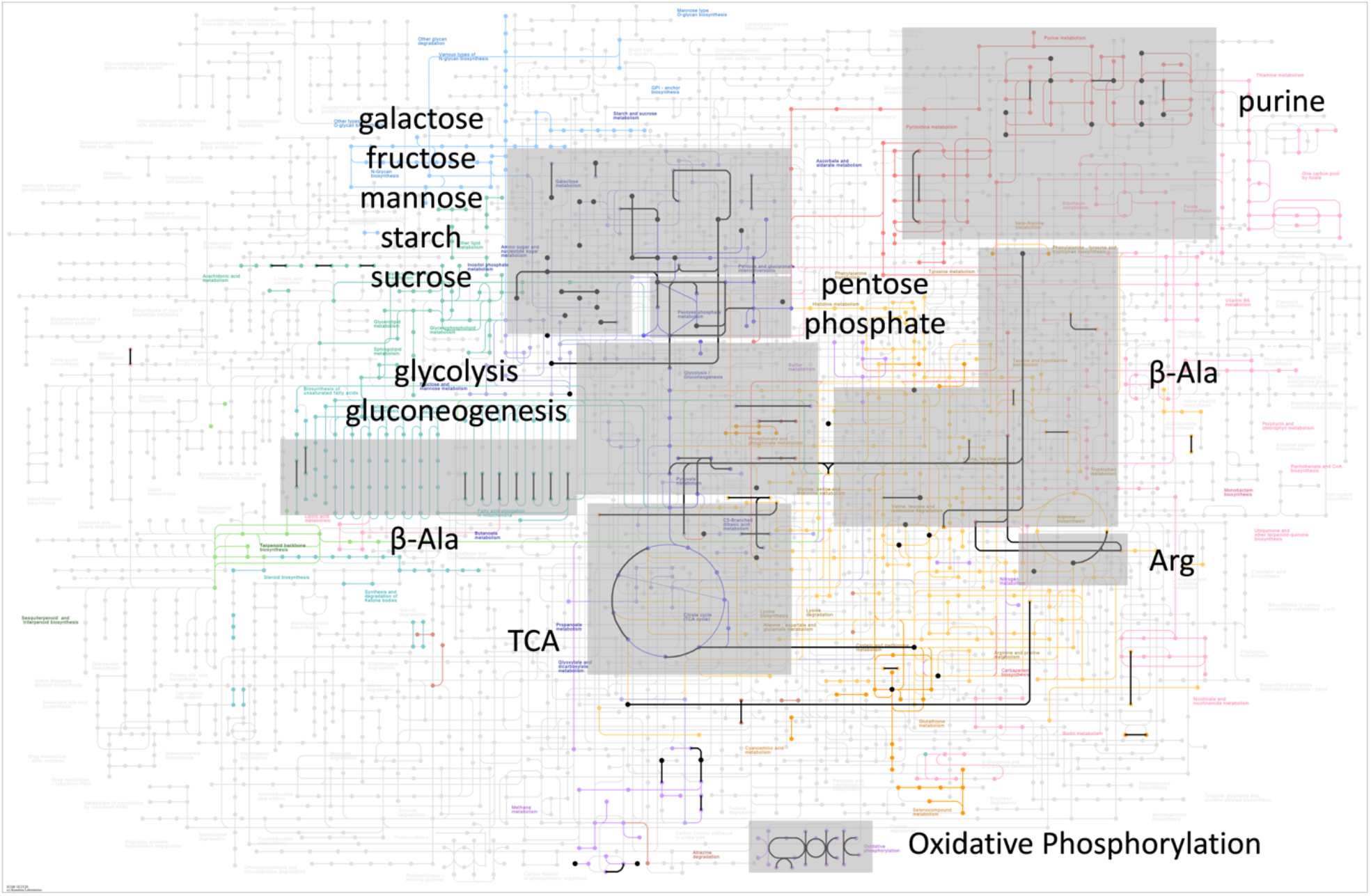
Overview of DEGs and SPMs regulated by *GPA2* as determined by concentration analysis. KEGG Metabolic Pathway with DEGs and SPMs used as input for *gpa2* integration analysis, highlighted using black lines and dots, respectively. Grey box is drawn over clusters associated with a specific pathway. Pathway names are color coded as detailed in Figure 3.

**Figure 6.**
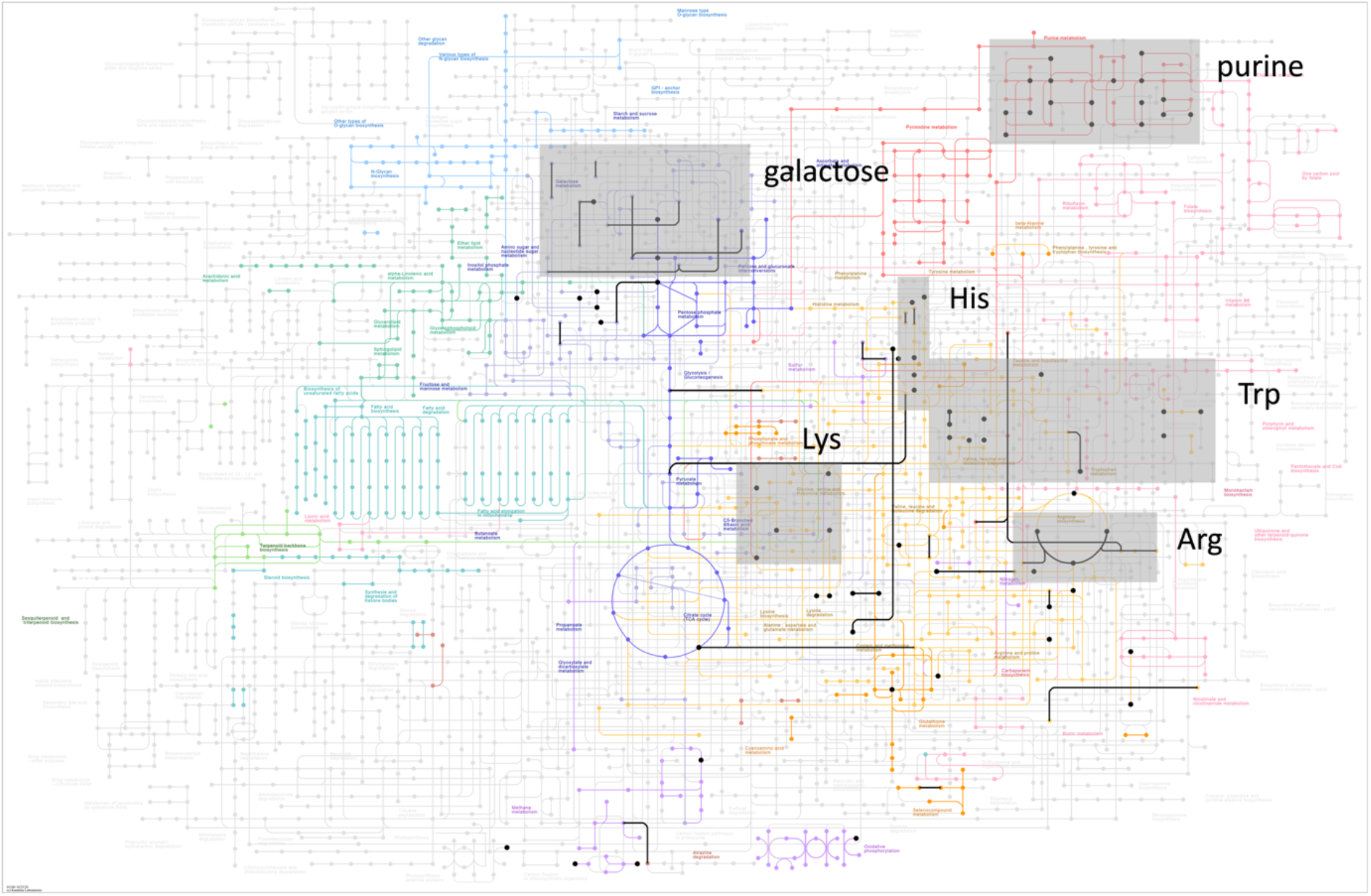
Overview of DEGs and SPMs regulated by *ASC1* as determined by concentration analysis. KEGG Metabolic Pathway with DEGs and SPMs used as input for *asc1* integration analysis, highlighted using black lines and dots, respectively. Grey box is drawn over clusters associated with a specific pathway. Pathway names are color coded as detailed in Figure 3.

In summary, our concentration analysis provides new and complementary information about glucose signaling. In particular, using an integrated transcriptomics and metabolomics approach (39, 40), we were able to confirm and consolidate changes seen at the metabolic or transcriptional level. For example, the integration analysis confirmed results obtained at the single-omics level; that Gpa2 affects pathways related to sugar utilization while Asc1 affects pathways related to amino acid metabolism. Second, integration analysis revealed information that might have been hidden using single-omics analysis methods. Only when the two datasets were combined did several important pathways meet the threshold of significance. For example, when comparing *gpa2* to wildtype, integration analysis established a substantial and statistically significant role for the Gα protein Gpa2 in glycolysis and gluconeogenesis. Of the fourteen features that emerged from our integrated analysis of this pathway, seven came from metabolomics and seven came from transcriptomics, neither of which met the threshold of significance on its own (Table 2 and Table S1, Supporting Information). Likewise, when comparing *asc1* to wildtype, integration analysis helped us to establish a specific, substantial and statistically significant role for the Gβ protein Asc1 in cysteine and methionine metabolism (Table 3 and Table S2, Supporting Information). Since the metabolic pathway is comprised of enzymes (gene products) and metabolites (enzyme products) it is noteworthy that both are regulated in a similar manner, even if the numbers obtained from each analytical method is small. Finally, our integration analysis allowed us to narrow the role for Gpa2 from very general effects on sugar metabolism to a more specific role in regulating glucose and ATP. Conversely, our integration analysis allowed us to show a broader role for Asc1 in amino acid metabolism, one not limited to a specific small subset of amino acids.

### Sensitivity analysis

The previous section compares the role of each signaling component in establishing transcript and metabolite concentration after glucose addition. Another way to explore the consequences of glucose sensing is to instead measure changes in response amplitude (sensitivity analysis), defined here as the difference between mutant (high minus low glucose, mutantH-mutantL) and wildtype (high minus low glucose, wtH-wtL). To put this in a biological context, a change in response amplitude reflects the role of a given component (Asc1 or Gpa2) in regulating the *relative* level of metabolites and/or genes, in response to a perturbation. A case in point is the relative change in cAMP after glucose addition, where the fold-change in its abundance is detected by the cell, regardless of the starting concentration. Differences detected using this approach are illustrated in Figure 2B.

We first compared transcriptional changes in wildtype and mutant cells, using the interaction term in DESeq2 (41). In this analysis we found altered sensitivity for 996 and 617 genes (s-DEGs) when comparing *gpa2* with wildtype and comparing *asc1* with wildtype, respectively. In sensitivity analysis we defined s-DEGs as having altered sensitivity changes with an adjusted p-value <0.05 and absolute fold-change value >1. Using GSEA we determined that Gpa2 regulates the sensitivity of transcripts linked to ten pathways including N-glycan biosynthesis, sugar-related metabolism and steroid biosynthesis (Table 4). These changes are likely related to cell membrane and cell wall biosynthesis, leading to cell division. Asc1 regulates twelve pathways, including ribosome biogenesis and sugar-related metabolism (Table 5). A Venn diagram for these comparisons reveals that both *gpa2* and *asc1* had similar effects on 47 s-DEGs, of which 22 were up-regulated and 25 down-regulated (Figure 7A and Table S7, Supporting Information). These shared s-DEGs were enriched for arginine and purine metabolism (Figure 7B). For purine metabolism, the mutants had opposing effects for the concentration analysis (as discussed earlier) but correspondent effects for the sensitivity analysis. This suggests that purine metabolism is an important target of this glucose-sensing pathway and is congruent with the role of glucose as a precursor in purine biosynthesis. The individual mutants were over-represented for four pathways related to ribosome biogenesis, monobactam biosynthesis and RNA polymerase (*asc1*, Figure 7C and Table S7, Supporting Information) and six pathways linked to amino acid and glycerophospholipid metabolism (*gpa2*, Figure 7D and Table S7, Supporting Information). These results reveal that Gpa2 uniquely affects the sensitivity of genes related to glycan and lipid metabolism (processes related to cell membrane and cell wall synthesis), as well as some cofactors related to amino acids, while Asc1 mainly affects the sensitivity of genes related to ribosome and translation (processes related to new protein synthesis), consistent with its role as a subunit of the 40S ribosome.

**Table 4.** Single- and Multi-omics integration results for gpa2 by sensitivity analysis. First block shows GSEA for transcriptomics with adjusted p-value <0.05, arranged in ascending order; second block shows MetaboAnalystR pathway enrichment analysis for metabolomics with combined p-value <0.05, arranged in ascending order; third block shows MetaboAnalystR joint pathway analysis with adjusted p-value <0.05, arranged in ascending order, as detailed in Methods.

| Transcriptomics |  | Metabolomics |  | Integration |  |
| --- | --- | --- | --- | --- | --- |
| enriched pathways | adjusted p-value | enriched pathways | combined p-value | enriched pathways | adjusted p-value |
| N-Glycan biosynthesis | 0.0050 | Fructose and mannose metabolism | 0.0044 | Glycolysis or Gluconeogenesis | 2.32E-06 |
| Various types of N-glycan biosynthesis | 0.0050 | Glycolysis / Gluconeogenesis | 0.0183 | Galactose metabolism | 2.32E-06 |
| DNA replication | 0.0050 | Methane metabolism | 0.0211 | Cell cycle | 1.61E-05 |
| Aminoacyl-tRNA biosynthesis | 0.0050 | Porphyrin and chlorophyll metabolism | 0.0473 | Pentose phosphate pathway | 2.03E-05 |
| Cysteine and methionine metabolism | 0.0050 |  |  | Starch and sucrose metabolism | 3.41E-05 |
| Starch and sucrose metabolism | 0.0050 |  |  | MAPK signaling pathway | 0.0002 |
| Galactose metabolism | 0.0050 |  |  | Meiosis | 0.0010 |
| Cell cycle | 0.0056 |  |  | Fructose and mannose metabolism | 0.0010 |
| Mismatch repair | 0.0063 |  |  | Purine metabolism | 0.0025 |
| Steroid biosynthesis | 0.0308 |  |  | Cysteine and methionine metabolism | 0.0025 |
|  |  |  |  | Protein processing in endoplasmic reticulum | 0.0096 |
|  |  |  |  | Peroxisome | 0.0124 |
|  |  |  |  | Spliceosome | 0.0230 |
|  |  |  |  | Pyruvate metabolism | 0.0238 |
|  |  |  |  | DNA replication | 0.0238 |
|  |  |  |  | Longevity regulating pathway | 0.0238 |
|  |  |  |  | Glycerolipid metabolism | 0.0238 |
|  |  |  |  | beta-Alanine metabolism | 0.0238 |
|  |  |  |  | Steroid biosynthesis | 0.0238 |
|  |  |  |  | Amino sugar and nucleotide sugar metabolism | 0.0238 |
|  |  |  |  | Alanine, aspartate and glutamate metabolism | 0.0238 |
|  |  |  |  | Inositol phosphate metabolism | 0.0241 |

**Table 5.** Single- and Multi-omics integration results for asc1 by sensitivity analysis. First block shows GSEA for transcriptomics with adjusted p-value <0.05, arranged in ascending order; second block shows MetaboAnalystR pathway enrichment analysis for metabolomics with combined p-value <0.05, arranged in ascending order; third block shows MetaboAnalystR joint pathway analysis with adjusted p-value <0.05, arranged in ascending order, as detailed in Methods.

| Transcriptomics |  | Metabolomics |  | Integration |  |
| --- | --- | --- | --- | --- | --- |
| enriched pathways | adjusted p-value | enriched pathways | combined p-value | enriched pathways | adjusted p-value |
| Ribosome | 0.0071 | Lysine degradation | 0.0082 | Ribosome | 4.76E-98 |
| Ribosome biogenesis in eukaryotes | 0.0071 | Cysteine and methionine metabolism | 0.0473 | Ribosome biogenesis in eukaryotes | 1.87E-21 |
| Glyoxylate and dicarboxylate metabolism | 0.0071 |  |  | RNA polymerase | 0.0015 |
| RNA polymerase | 0.0071 |  |  | Galactose metabolism | 0.0015 |
| Peroxisome | 0.0071 |  |  | Pentose phosphate pathway | 0.0422 |
| Galactose metabolism | 0.0104 |  |  | Purine metabolism | 0.0422 |
| Citrate cycle (TCA cycle) | 0.0141 |  |  | Arginine biosynthesis | 0.0422 |
| RNA transport | 0.0196 |  |  |  |  |
| Starch and sucrose metabolism | 0.0241 |  |  |  |  |
| Carbon metabolism | 0.0479 |  |  |  |  |
| Propanoate metabolism | 0.0479 |  |  |  |  |
| Aminoacyl-tRNA biosynthesis | 0.0498 |  |  |  |  |

**Figure 7.**
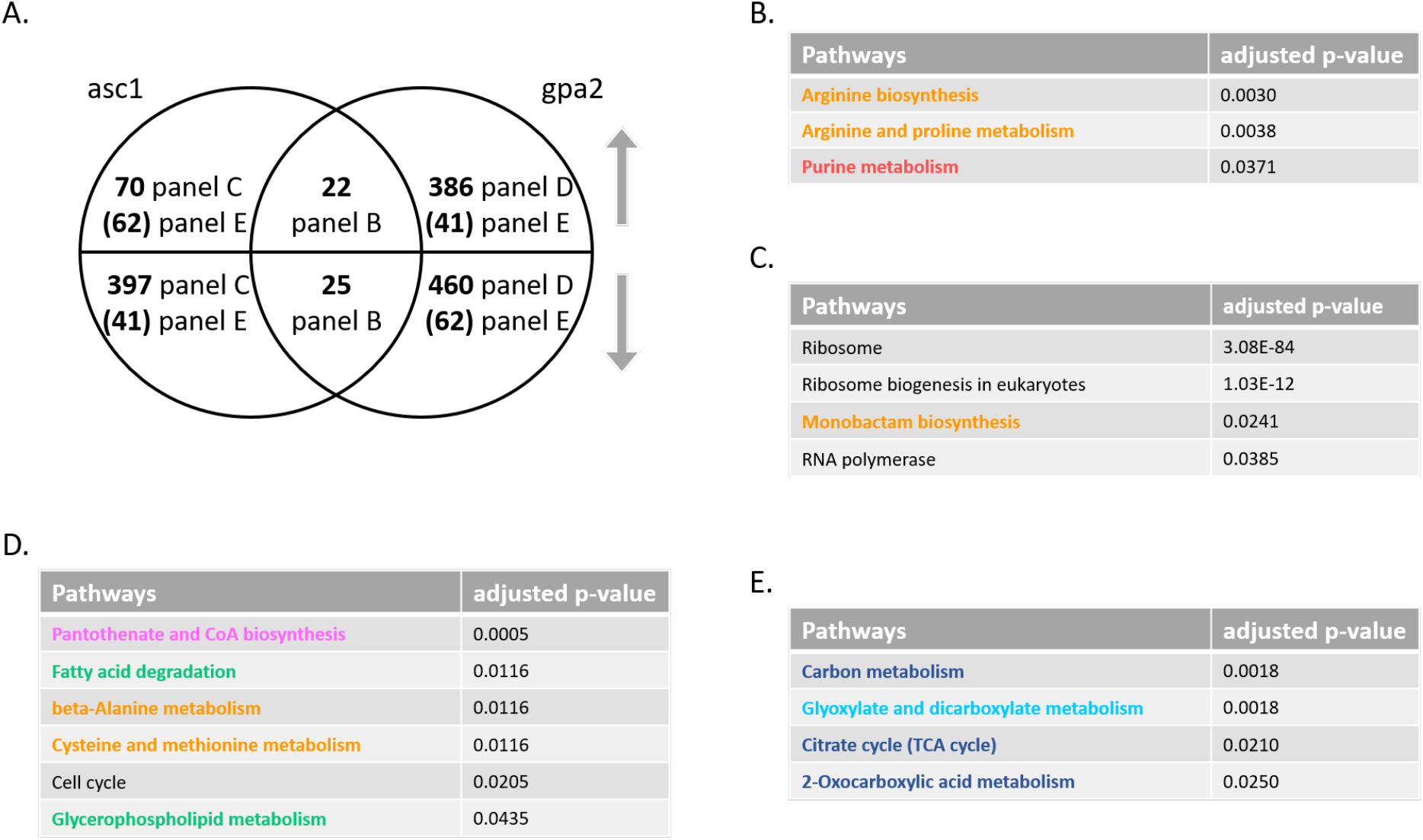
Sensitivity analysis of differentially expressed genes (s-DEGs) after glucose addition. A) Venn diagram of subsets of s-DEGs, before and after glucose addition for each mutant (mutantH-mutantL) and wildtype (wtH-wtL). Upper semicircle shows up-regulated s-DEGs and lower semicircle shows down-regulated s-DEGs. Numbers in parenthesis are shared s-DEGs regulated in the opposite direction, placed in the area corresponding to the direction of regulation. subset of s-DEGs used for ORA analysis that are B) shared and change in the same direction; C) unique to *asc1*; D) unique to *gpa2*; E) shared and change in the opposite direction. Pathway names are color coded according to KEGG metabolic pathway map as detailed in Figure 3. Pathways with adjusted p-value <0.05 is listed with ascending order in the table.

While each of the mutants had unique effects, there were an additional 103 s-DEGs regulated by both mutants but in opposite directions (Table S7, Supporting Information). Whereas concentration analysis revealed opposing effects for a small group of s-DEGs related to transposon elements, our sensitivity analysis revealed opposing effects for s-DEGs related to sugar metabolism (Figure 7E). For the 62 s-DEGs that were up-regulated in *asc1* and down-regulated in *gpa2*, most were due to a change in basal expression (at low glucose) (Figure S8, Supporting Information). For the remaining 41 s-DEGs (Figure 7E), the pattern was more complex, with mutants affecting basal expression or induced expression or both (Figure S9, Supporting Information).

We then performed generalized linear modeling to detect significant differences in metabolite/peak levels, comparing (wtH-wtL) and (mutantH-mutantL), followed by pathway enrichment analysis in MetaboAnalystR, as detailed above. By this approach we determined that the *gpa2* strain was enriched for four pathways, primarily sugar-related, while *asc1* was enriched for two pathways including lysine, cysteine, and methionine metabolism (Tables 4 and 5). A Venn diagram shows that both *gpa2* and *asc1* had similar effects on 16 s-SPMs (5 increased, 11 decreased, defined as those with an adjusted p-value <0.05), with no pathway over-represented (Figure 8A and Table S8, Supporting Information). *asc1* uniquely affected 27 s-SPMs, with diverse functions, while *gpa2* uniquely affected 31 s-SPMs (18 increased, 13 decreased), enriched for steroid biosynthesis (Figure 8B and Table S8, Supporting Information). These results confirm the role of Gpa2 in lipid metabolism, as shown above. Most notably, the mutants had opposing effects on the sensitivity of an additional 42 compounds primarily related to glucose metabolism, including glycolysis and gluconeogenesis, amino and nucleotide sugars, as well as the metabolism of galactose, fructose, mannose, sucrose and starch (Figure 8C and Table S8, Supporting Information).

**Figure 8.**
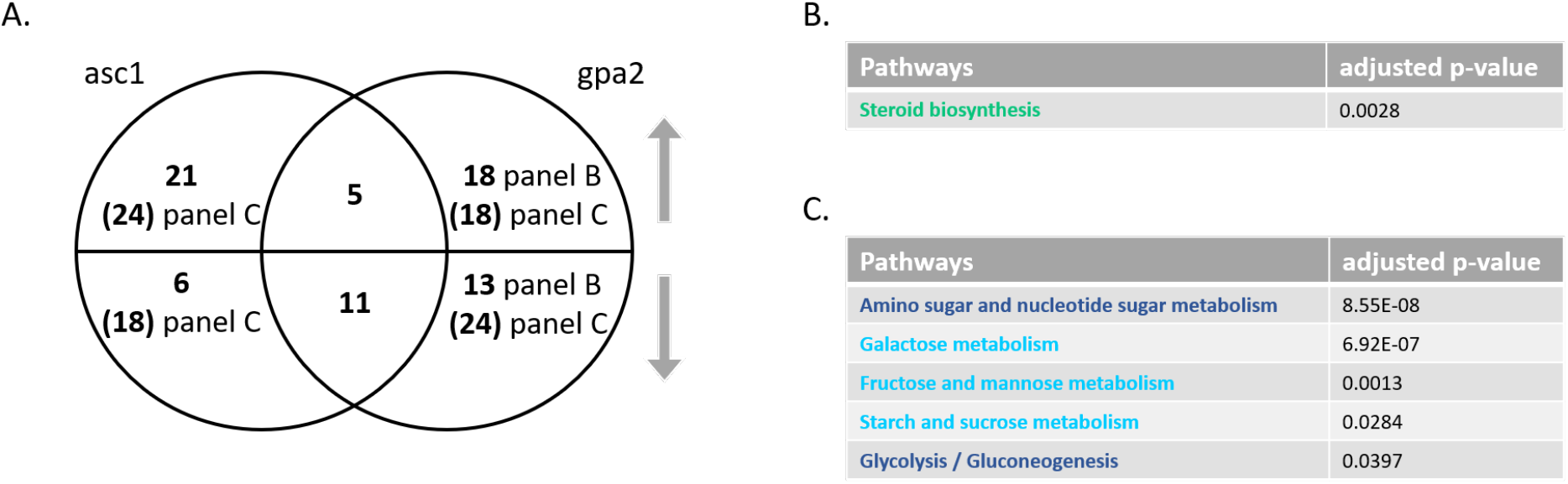
Sensitivity analysis of significantly perturbed metabolites (s-SPMs) after glucose addition. A) Venn diagram of subsets of s-SPMs, before and after glucose addition for each mutant (mutantH-mutantL) and wildtype (wtH-wtL). Upper semicircle shows up-regulated s-SPMs and lower semicircle shows down-regulated s-SPMs. Numbers in parenthesis are shared s-SPMs regulated in the opposite direction, placed in the area corresponding to the direction of regulation. subset of s-SPMs used for ORA analysis that are B) unique to *gpa2*; C) shared and changed in the opposite direction. Shared s-SPMs changed in the same direction and s-SPMs unique to *asc1* are not over-represented in any pathway and are therefore not shown. Pathway names are color coded according to KEGG metabolic pathway map as detailed in Figure 3. Pathways with adjusted p-value <0.05 is listed with ascending order in the table.

The results above reveal important differences between the two G protein subunits, as determined by concentration and sensitivity analysis. In general, the two mutants responded in opposing ways to glucose addition, and did so at both the metabolic and transcriptional levels. Across all gene level comparisons, only arginine metabolism was affected similarly by the two mutants (Figures 3 and 7). Given the opposing roles of Asc1 and Gpa2 in other processes it is noteworthy that aspects of arginine metabolism were also affected, and in the same direction, by a mutant lacking the shared activator Gpr1 (Tables S5 and S9, Supporting Information). More specifically, all three mutants led to the induction of five genes (*AGP1, MEP1, DAL2, GDH1*, and *EDS1*). Quantifiable changes in these gene transcripts may prove useful as reporters of glucose-mediated GPCR signaling.

We then applied integration analysis and found that *gpa2* was enriched in 22 pathways while *asc1* was enriched in seven pathways (Tables 4 and 5). By our sensitivity analysis, we found that both mutants affected genes or metabolites involved in the pentose phosphate pathway, as well as purine and galactose metabolism. The *gpa2* strain uniquely regulated 19 pathways, involved in diverse cellular functions (Table 4), while *asc1* uniquely regulated 4 pathways, all related to its known role in ribosome function as well as arginine metabolism (Table 5). Thus, the relationship of Asc1 to amino acids and ribosome function is reflected in both concentration and sensitivity analysis. By concentration analysis, the effects of *gpa2* were primarily related to sugar metabolism, while sensitivity analysis revealed a greater diversity of processes including amino acids and lipids, in addition to sugars. This indicates that while Gpa2 is crucial in maintaining sugar metabolism, it also interacts with genes and metabolites related to other non-sugar species to ensure their proper response amplitude upon sugar addition. Taken together, it is evident that concentration and sensitivity analysis provide information that is complementary and biologically meaningful.

Finally, in order to visualize the functional relationship of Gpa2 and Asc1, we projected the inputs of our integration analysis onto the comprehensive yeast Metabolic Pathways diagram provided in KEGG (Figures 9 and 10). When comparing the two plots, it is evident that Gpa2 affects the sensitivity of a larger number of genes and metabolites, as compared with Asc1. The effects of *gpa2* were pervasive and included glucose, other sugars, lipids, amino acids and purine nucleotides (Figure 9). In contrast, the effects of *asc1* centered on pentose phosphate, purine and arginine metabolism (Figure 10). In addition, and as anticipated given its role in ribosome assembly, *asc1* uniquely affected processes related to RNA polymerase and ribosome biogenesis, which are not included in the Metabolic KEGG map (Table 5). These findings reveal that, although the two G protein subunits each had substantial effects by concentration analysis, the Gα subunit had by far the largest effect on sensitivity analysis. Collectively, these data indicate that Gpa2 affects the amplitude of the response to glucose, and does so for an especially large group of genes and metabolites.

**Figure 9.**
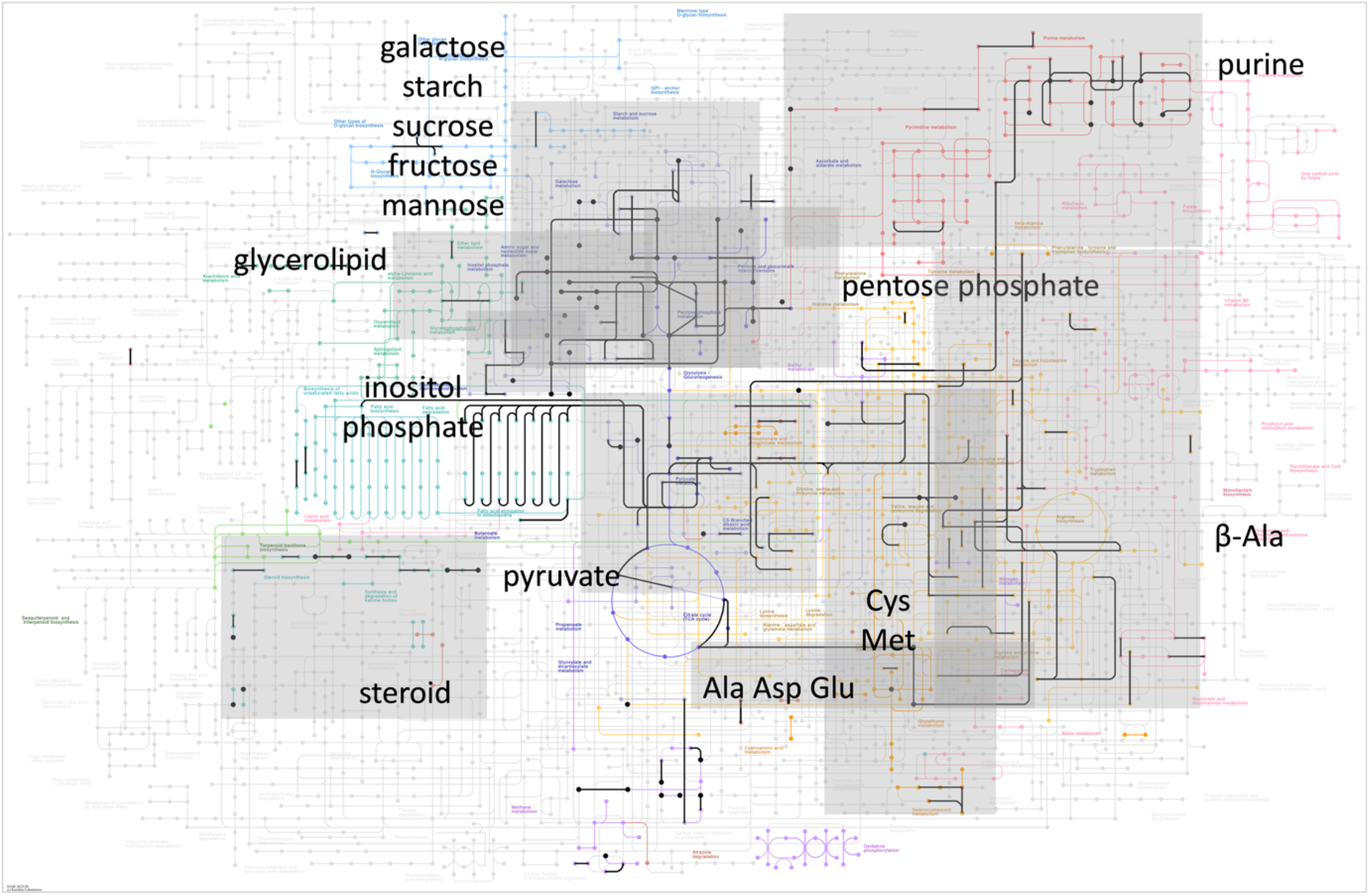
Overview of s-DEGs and s-SPMs regulated by GPA2 as determined by sensitivity analysis. KEGG Metabolic Pathway with s-DEGs and s-SPMs used as input for *gpa2* integration analysis, highlighted using black lines and dots, respectively. Grey box is drawn over clusters associated with a specific pathway. Pathway names are color coded as detailed in Figure 3.

**Figure 10.**
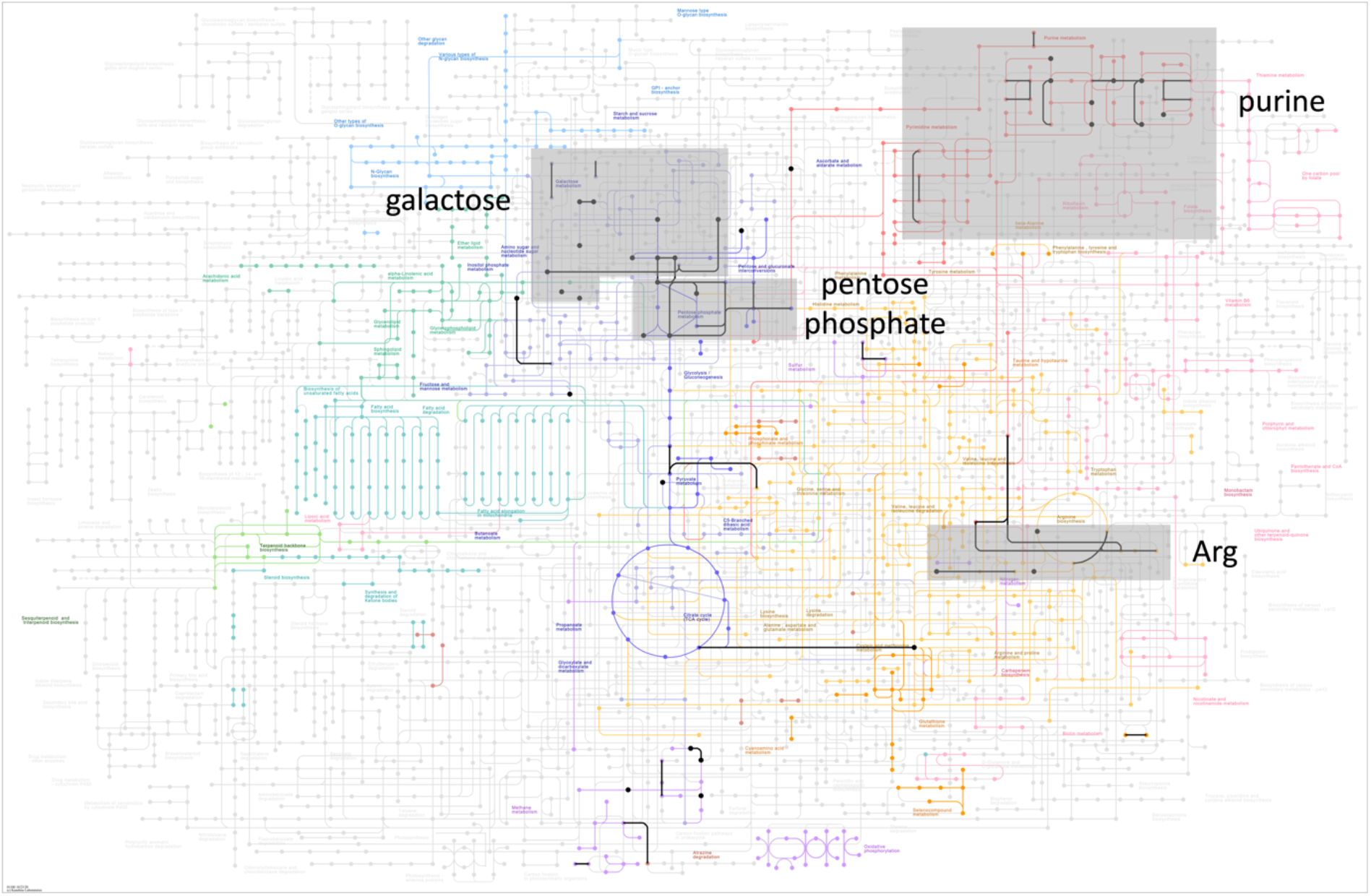
Overview of s-DEGs and s-SPMs regulated by ASC1 as determined by sensitivity analysis. KEGG Metabolic Pathway with s-DEGs and s-SPMs used as input for *asc1* integration analysis, highlighted using black lines and dots, respectively. Grey box is drawn over clusters associated with a specific pathway. Pathway names are color coded as detailed in Figure 3.

**Figure 11.**
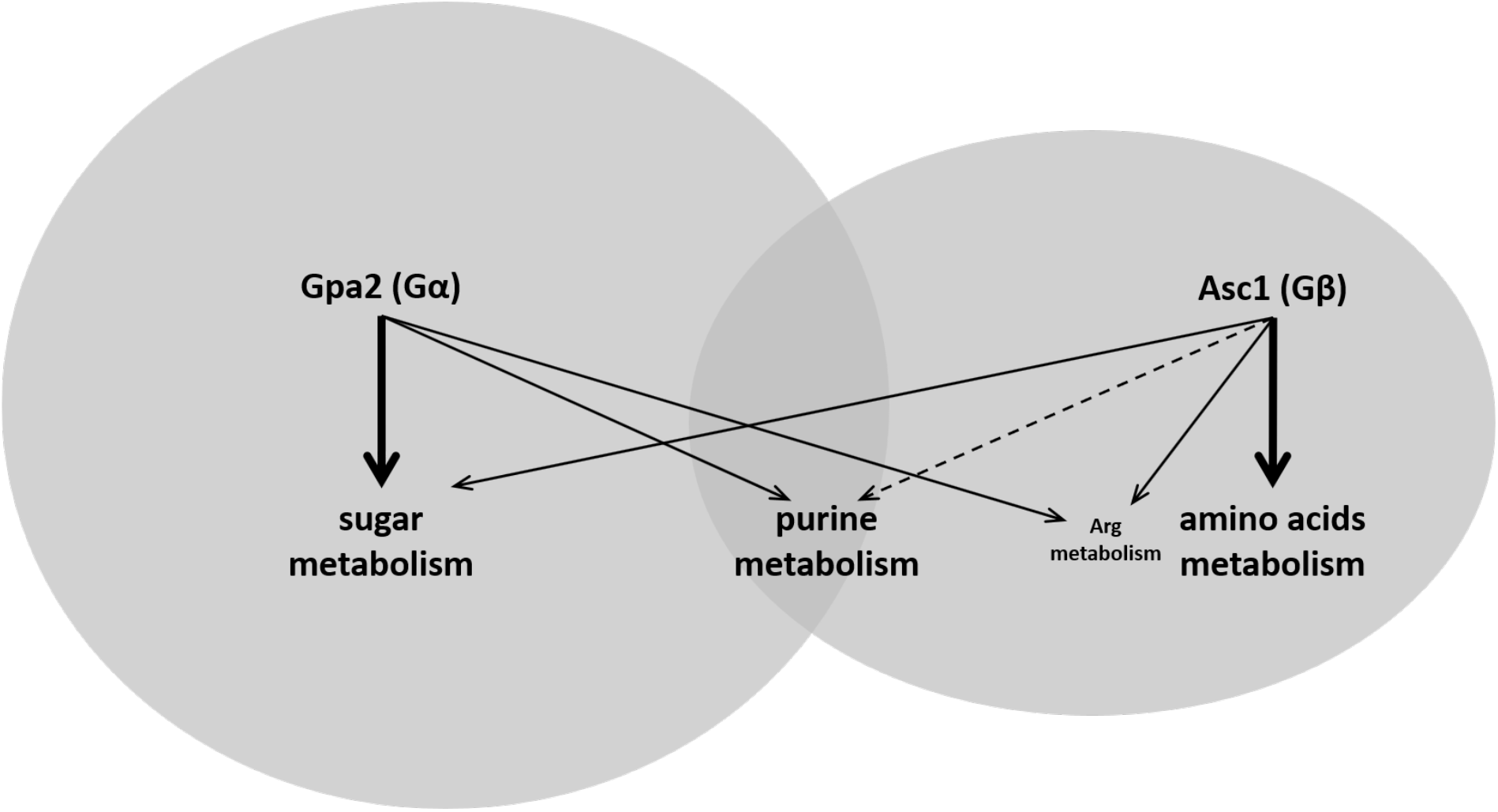
Summary of interactions. Gpa2 is primarily involved in regulating sugar metabolism while Asc1 is primarily involved in amino acid metabolism, as indicated with thick lines. Gpa2 and Asc1 have shared effects on arginine metabolism and opposing effects on purine metabolism, as indicated with solid and dashed lines. Gpa2 regulates a greater number of gene transcripts and was particularly important in determining the amplitude of response to glucose addition, as indicated with the larger grey sphere.

## DISCUSSION

The yeast *S. cerevisiae* has two functionally distinct GPCRs and two Gα proteins, but only a single canonical Gβγ. Here we considered the role of a second, “atypical”, Gβ protein Asc1 and compared its function with that of the cognate Gα protein Gpa2. Like other Gβ proteins, Asc1 binds preferentially to Gα-GDP, slows GDP-GTP exchange (9) and has a seven-bladed propeller domain structure (22, 42, 43). Like Gpa2, Asc1 binds directly to the effector adenylyl cyclase. Whereas Gpa2 stimulates the production of cAMP however, Asc1 has the opposite effect (9, 11). The effects on cAMP mirror some of the effects reported here, where Gpa2 and Asc1 have antagonistic effects on the purine metabolic pathway. It is possible that some of the documented effects on cAMP are the cause or consequence of changes to ATP biosynthesis.

While Asc1 and Gpa2 have opposing effects on cAMP production they have similar effects on haploid invasive growth (9, 33, 34), a process by which the cells form long branching filaments and exhibit increased adherence and invasion of the substratum. This growth phenotype occurs during periods of glucose limitation (35), possibly in an attempt to direct colony expansion to sites of greater nutrient availability. The fact that the Gα and Gβ have opposing effects on adenylyl cyclase but similar effects on invasive growth suggested to us that these proteins regulate processes other than adenylyl cyclase activation. To gain a better understanding of their relative contributions to cell physiology, we undertook an integrated metabolomics and transcriptomics analysis, comparing mutants that lack the Gα, or Gβ subunit. In this way, we established the existence of multiple shared and opposing outputs. While much remains to be done, we anticipate that the glucose-dependent metabolic and transcriptional changes reported here will provide some clues as to the underlying biochemical processes responsible for the signaling events mediated by these two G protein subunits.

As noted above, Gpa2 and Asc1 have opposing effects on adenylyl cyclase, as well as other metabolic processes in the cell. Such competing interactions mirror those that occur in animal cells, where Gα and Gβγ proteins often have antagonistic effects on adenylyl cyclase activity (12). While seemingly paradoxical, this design has several useful properties (44). First, such “antagonistic bifunctionality” results in an input-output relationship that is insensitive to naturally occurring fluctuations in protein abundance (“robustness”). Since Gα does not exist without Gβ, an increase in the positive regulator leads to a concomitant increase in the negative regulator. Without such bifunctionality the positive or negative output may lead to spurious signaling (“noise”) by one or the other subunit. Second, if the positive and negative regulators operate on different time scales they can produce outputs that are inherently transient. This might occur if there is a delay between the inactivation of Gα (following GTP hydrolysis) and the inactivation of Gβγ (following reassociation with Gα). Third, a built-in delay can confer a property known as fold-change detection. In this scenario, the amplitude of the output is proportional to the relative difference in, rather than the absolute concentration of, the input. This can occur when the negative regulator restores the output to the pre-stimulus baseline despite sustained activation by the positive regulator. Indeed, our fold-change analysis, or sensitivity analysis, allowed us to detect phenotypes that were not evident by concentration analysis alone. All of these features are characteristic of other G protein signaling systems. A familiar and well documented example is light detection by the GPCR rhodopsin; this system exhibits exceptionally low noise, a transient neural output, and a wide-dynamic range of light detection (45).

While many of the effects of Asc1 overlap with those of Gpa2, others appear to be unrelated to glucose sensing. We infer that these alternative outputs are related to the role of Asc1 as an accessory component of the ribosome, where it is required for some protein-protein interactions within the complex and for proper protein translation (18-23, 26). Other functions for Asc1 may exist. One possibility is that the effects of glucose on cell metabolism are somehow coordinated with changes in protein translation. That is to say, the protein translation machinery is made aware of changes in nutrient abundance, through the coordinated action of a shared protein Asc1. Indeed Asc1 appears to associate with the ribosome only under glucose-rich conditions (23), and regulates a subset of transcripts related to glycolysis, respiration, oxidative stress and glucose fermentation (26).

An alternative hypothesis is that the multiple functions of Asc1 are truly independent, a phenomenon for which there is ample precedent. The recruitment of a protein that originally evolved with one function to serve a second, unrelated function is an example of Darwinian exaptation, more colloquially known as “protein moonlighting” because it is analogous to a single person with two different jobs (46). This term is not applied to multi-domain proteins or to the products of alternative splicing or gene duplication. The first description of a moonlighting protein was argininosuccinate lyase, which has a well-established enzymatic function but which also serves as a structural protein responsible for the transparency of the lens and cornea (47, 48). Like Asc1, a large fraction of other proteins involved in glucose metabolism exhibit moonlighting behaviors; this includes the majority of enzymes involved in glycolysis and the tricarboxylic acid cycle (49, 50). There is also evidence for moonlighting by typical G proteins. The mammalian Gβ_2_ binds to Gα and G*γ* subunits but also assembles with the ubiquitin ligase DDB1-CUL4A-ROC1, where it helps to recruit the receptor kinase GRK2 for targeted degradation (51). Our identification of Asc1 as the Gβ for Gpa2 is like that of many other moonlighting proteins that have been identified because of their presence in unexpected multiprotein complexes or locations.

While it has most of the functions of a typical Gβ, Asc1 has a structure that makes it unique among Gβ protein subtypes (22, 42, 43). This too has precedent. Whereas Gpa1 binds to a typical Gβ (Ste4), which is necessary for pheromone-induced mating, it also associates with an atypical Gβ called Vps15, which promotes autophagy through activation of phosphatidylinositol 3-kinase (4, 52). Our x-ray structure determination revealed that Vps15 has a 7-bladed propeller domain structure, similar to that of other Gβ proteins, as well as a protein kinase domain of unknown function. The discovery of atypical and multifunctional Gβ proteins, including Asc1 and Vps15, suggests that the superfamily of Gβ subunits may be far larger and more complex than previously recognized (53).

It has long been appreciated that the glucose-sensing pathway in yeast employs a G protein-coupled receptor, and that activation of this pathway suppresses invasive growth. Despite the nutritional and societal importance of glucose fermentation in yeast however, the mechanisms and molecular consequences of glucose sensing have remained obscure. Through an analysis of individual gene deletion mutants, and by integrating transcriptomic and metabolomic measurements, we have determined the contributions of the Gα and Gβ protein subunits to transcription and metabolism. We anticipate that the approaches implemented here will be useful for investigating other GPCR pathways and other moonlighting proteins, including G proteins, that participate in multiple, seemingly unrelated, biological processes.

## METHODS

### Yeast strains

The prototrophic (wildtype) strain used throughout was constructed from BY4741 (MAT**a** *his3*Δ1 *leu2*Δ0 *met15*Δ0 *ura3*Δ0). *HIS3, LEU2, MET15* and *URA3* were integrated at the endogenous loci with sequence amplified by PCR from S288C strain DNA. All single mutants (*gpr1, gpa2, asc1*) were constructed by transforming the wildtype strain with corresponding sequence from the Yeast Knock-Out collection that replaces the target gene with KanMX4 (54).

### Cell preparation

All strains were cultivated in the same way and maintained at 30 °C unless otherwise indicated. Cells were inoculated into Synthetic Complete (SC) (2% glucose) overnight and grown to saturation, then back diluted and kept in log phase overnight. The next morning, the culture was harvested when OD reached 1.0. For each genotype, 90 ml of culture (OD=1.0) was split into 2 tubes (45 ml each) for later high or low glucose treatment. For each tube, cells were centrifuged and resuspended twice with SC (0.05% glucose). Cells were then resuspended into 10 ml SC (0.05% glucose) and cultivated for 1 h.

For high glucose treatment, 245 μL 65.5% glucose was added to 10 mL cell culture and for low glucose treatment, 245 μL 0.05% glucose was added to 10 mL cell culture, each for exactly 2 min (metabolomics) or 10 min (transcriptomics). The 10 min time point was selected based on a pilot time course experiment (2, 5, 10, 15, 20, 30 and 45 min), showing that this is the earliest time point that captures most of the transcripts affected by the glucose treatment.

For metabolomics, each replicate consisting of 3 mL of cell culture was mixed with 45 mL cold pure methanol on dry ice. After 5 min, cells were centrifuged in a precooled rotor (−80 °C). After discarding the supernatant, cell pellets were immediately stored at −80 °C. A small aliquot of each sample was saved to manually determine cell density with a hemocytometer.

For RNA-seq, 500 μL of cell culture was aliquoted into a 1.7 mL microfuge tube and centrifuged at 1000 x *g* for 1 min at 4 °C. After discarding supernatant, the cell pellet was flash frozen by liquid nitrogen and stored at −80 °C.

### Sample preparation for RNA-seq

Cell pellets stored at −80 °C were resuspended with 600 μL buffer RLT 1% (v/v) 2-mercaptoethanol from the QIAGEN RNeasy Mini Kit (Cat No.: 74106) and then added into 2 mL OMNI prefilled ceramic bead tubes (SKU: 19-632). Tubes were loaded onto an OMNI Bead Mill Homogenizer (SKU:19-040E) for 3 beating cycles. For each cycle, samples were agitated at 5 m/s for 1 min at 4 °C and then cooled on ice for 3 min between each cycle. The resulting lysate was clarified by centrifugation at 11,000 xg and then used for total RNA extraction with QIAGEN RNeasy Mini Kit (Cat No.: 74106) with on-column DNase digestion according to manufacturer’s instructions. Extracted total RNA for each sample was checked for purity and quantified with Invitrogen Qubit 2.0 Fluorometer (Cat No.: Q32866) and Qubit RNA HS Assay kit (Cat No.: Q32855) according to manufacturer’s instructions.

### Sample preparation for metabolomics

Frozen cell pellets were resuspended with extraction reagent (8:2 methanol-water solution) to 3×10^8^ cell/mL and then transferred into 2 mL ceramic bead MagNalyser tubes. Blank samples were prepared by adding 1300 μL of extraction reagent with no cells to a MagNalyser tube with ceramic beads. Tubes were subjected to homogenization, with Bead Ruptor Elite Bead Mill Homogenizer (OMNI International) at 6.0 m/s for 40 sec in 2 cycles at room temperature. This step was repeated twice. All samples were then centrifuged at 16,000 xg for 10 min at 4 °C. 500 μL of the supernatant was transferred into low-bind 1.7 mL microfuge tubes. Total pools were made by combining an additional 65 μL of the supernatant from each sample and then aliquoting this mixture into low-bind 1.7 mL tubes at a volume of 500 μL. The remaining supernatant was stored at −80 °C for repeat experiments if necessary. For all experimental samples, pooled samples and blanks were dried using a speedvac vacuum concentrator overnight. Dried samples were stored at −80 °C.

Before LC-MS analysis, 100 μL of reconstitution buffer (95:5 water:methanol with 500 ng/mL tryptophan d-5) was added to each dried sample. All tubes were vortexed at 5000 rpm for 10 min and then centrifuged at room temperature at 16,000 xg for 4 min. Supernatant was transferred into autosampler vials for LC-MS.

### RNA library preparation

RNA libraries were prepared with Kapa stranded mRNA-seq kits, with KAPA mRNA Capture Beads (KAPA code: KK8421; Roche Cat No.: 07962207001) through the UNC High Throughput Sequencing Facility. All procedures were according to manufacturer’s instructions.

### RNA Sequence analysis

Quality of raw sequence was checked with the FASTQC algorithm (http://www.bioinformatics.babraham.ac.uk/projects/fastqc/). Sequence alignment to genome indices, generated based on *Saccharomyces cerevisiae* data downloaded from Ensembl.org, was performed with the STAR algorithm (55). Quantification on a transcriptome level was performed with the SALMON algorithm (56). The quantified data were then analyzed with the DESeq2 package in R (57), which provides a means to determine differences in transcript abundance using a negative binomial generalized linear model (41). Differentially Expressed Genes (DEGs for concentration analysis, s-DEGs for sensitivity analysis) are defined as having adjusted p-value <0.05 and absolute fold-change > 1.

PCA analysis was performed using internal PCA function of DESeq2 package with variance stabilizing transformation (vst) normalized data.

For concentration analysis, sequencing results for wildtype and all mutants after glucose addition were used as input and were analyzed with the design formula = ~batch+genotype. Here ‘batch’ is incorporated to account for batch effects of sample preparation. For sensitivity analysis, sequencing results for wildtype and all mutants before and after glucose addition were used as input and were analyzed with the design formula = ~batch+genotype+treatment+genotype:treatment. Here ‘treatment’ is equivalent to glucose addition. The interaction term ‘genotype:treatment’ is included to estimate how the response amplitude of each mutant is different from wildtype, that is (mutantH-mutantL)-(wtH-wtL).

### UHPLC high-resolution Orbitrap MS metabolomics data acquisition

Metabolomics data were acquired on a Vanquish UHPLC system coupled to a QExactive™ HF-X Hybrid Quadrupole-Orbitrap Mass Spectrometer (Thermo Fisher Scientific, San Jose, CA), as described previously (58). Our UPLC–MS reversed phase platform was established based on published methods (59, 60). Metabolites were separated using an HSS T3 C18 column (2.1 × 100 mm, 1.7 μm, Waters Corporation) at 50 °C with binary mobile phase of water (A) and methanol (B), each containing 0.1% formic acid (v/v). The UHPLC linear gradient started from 2% B, and increased to 100% B in 16 min, then held for 4 min, with the flow rate at 400 μL/min. The untargeted data were acquired in positive mode from 70 to 1050 m/z using the data-dependent acquisition mode.

### Metabolomics data normalization and filtration

Progenesis QI (version 2.1, Waters Corporation) was used for peak picking, alignment, and normalization as described previously (58). Samples were randomized and run within two batches with blanks and pools interspersed at a rate of 10%. Starting from the un-normalized data for each batch runs, the data were filtered so as to only include signals with an average intensity fold change of 3.0 or greater in the total pools compared to the blanks. Individual samples (including pools, blanks, and study samples) were then normalized to a reference sample that was selected by Progenesis from the total pools via a function named “normalize to all”. Signals were then excluded that were significantly different between pools of batch 1 and pools of batch 2 based on an ANOVA comparison calculated in Progenesis (q <0.05). After normalization and filtration, 2397 signals passed the QC procedures and were used for further analysis.

The filtered and normalized data was mean-centered and Pareto scaled prior to conducting the unsupervised principal components analysis using the ropls R package

### In-house Compound identification and annotation

Peaks were identified or annotated by Progenesis QI through matching to an in-house experimental standards library generated by acquiring data for approximately 1000 compounds under conditions identical to study samples, as well as to public databases (including HMDB, METLIN and NIST), as described previously (58). Identifications and annotations were assigned using available data for retention time (RT), exact mass (MS), MS/MS fragmentation pattern, and isotopic ion pattern. The identification or annotation of each signal is provided in Supporting Information. Signals/metabolites that matched to the in-house experimental standards library by (a) RT, MS, and MS/MS are labeled as OL1, or (b) by RT and MS are labeled OL2a. An OL2b label was provided for signals that match by MS and MS/MS to the in-house library that were outside the retention time tolerance (± 0.5 min) for the standards run under identical conditions. Signals matched to public databases are labeled as PDa (MS and experimental MS/MS), PDb (MS and theoretical MS/MS), PDc (MS and isotopic similarity or adducts), and PDd (MS only) are also provided (Supporting Information).

### Transcriptomics pathway enrichment analysis and Over-representation analysis

Pathway enrichment analysis for transcriptomics data was performed with ClusterProfiler package in R (28); Log2 fold-change for each comparison (mutantH vs wtH for concentration analysis or mutantH-mutantL vs wtH-wtL for sensitivity analysis) was extracted from corresponding DESeq2 analysis. GSEA analysis was then performed with gseKEGG function, with organism set to ‘sce’ (*Saccharomyces cerevisiae*), permutation number set to 1000, minimal and maximal size for each analyzed geneset as 3 and 200, p-value cutoff set to 0.05, p-value adjustment method set to ‘BH’ (Benjamini-Hochberg). The KEGG sce database was used throughout, for both metabolomics and transcriptomics.

Over-representation analysis for the corresponding subsection of the Venn diagram was performed with the enrichKEGG function in ClusterProfiler package, with organism set to ‘sce’ (*Saccharomyces cerevisiae*), minimal and maximal size for each analyzed geneset as 3 and 200, p-value cutoff set to 0.05, p-value adjustment method set to ‘BH’ (Benjamini-Hochberg).

### Compound annotation, metabolic pathway enrichment analysis and Over-representation analysis

Compound annotation and pathway enrichment analysis for metabolomics was performed with the MetaboAnalystR 3.0 package in R (39, 40) https://www.metaboanalyst.ca/docs/RTutorial.xhtml). For compound annotations, molecular weight tolerance (ppm) was set to 3.0, analytical mode was set to positive and retention time was included. Pathway enrichment analysis was performed with ‘integ’ module (using both Mummichog V2.0 and GSEA) with the yeast KEGG database. The p-value threshold for Mummichog was set at 0.05.

For concentration analysis, normalized peak data, from Progenesis QI for wildtype and mutants after glucose addition, were used as input for MetaboAnalystR. For sensitivity analysis, normalized peak data for wildtype and mutants before and after glucose addition were used as inputs for generalized linear model:

AUC~genotype+treatment+genotype:treatment. AUC is the normalized area under curve for each peak. Treatment means before or after glucose addition. The interaction term estimated how the response amplitude of each mutant is different from wildtype, that is (mutantH-mutantL)-(wtH-wtL). The modeled p-value and t score for the interaction term associated with each peak were then used as inputs for pathway enrichment analysis.

Significantly perturbed metabolites (SPMs for concentration analysis, s-SPMs for sensitivity analysis) were defined as annotations that have adjusted p-value <0.05 (FDR) from the output of MetaboAnalystR. Significantly perturbed pathways were defined as having combined p-value <0.05 (Mummichog and GSEA).

Over-representation analysis for the corresponding subsection of the Venn diagram was performed with the Enrichment Analysis module in MetaboAnalystR, with KEGG ID for each metabolites as the input. FDR adjusted p-value <0.05 was the threshold for over-represented pathways.

### Integration of transcriptomics and metabolomics data

Integration analysis was performed with the ‘joint pathway analysis’ module of MetaboAnalystR (https://www.metaboanalyst.ca/docs/RTutorial.xhtml). For both concentration and sensitivity analysis, gene input together with log2 fold-change was generated based on the corresponding DESeq2 analysis, with the threshold set as adjusted p-value <0.05 and absolute log2 fold-change >1 (DEGs or s-DEGs); metabolite input together with log2 fold-change was generated based on MetaboAnalystR analysis, with the threshold set as adjusted p-value <0.05 (SPMs or s-SPMs). Integration analysis was performed on ‘all pathways’, which includes both metabolic pathways as well as gene-only pathways. Enrichment analysis was performed using ‘Hypergeometric test’. Topology measure was set to ‘Degree Centrality’. Integration method was set to ‘combine queries’, which is a tight integration method with genes and metabolites pooled into a single query and used to perform enrichment analysis within their “pooled universe”. Significantly enriched pathways were defined as having FDR adjusted p-value <0.05.

## ACKNOWLEDGEMENTS

We thank Daniel Dominguez and Patrick Maxwell for helpful discussion and comments on the manuscript. RNAseq was performed by Amy Perou and Piotr Mieczkowski at the High Throughput Sequencing Facility at UNC Chapel Hill. Metabolomics data capture, data analysis, and identification and annotation were conducted in the Susan Sumner Laboratory located at the Nutrition Research Institute at UNC Chapel Hill.

